# The combination of differential expression and differential network connectivity analyses identifies RNA splicing and processing as common pathways altered with age across human tissues

**DOI:** 10.1101/2023.05.26.542445

**Authors:** Caio M.P.F. Batalha, André Fujita, Nadja C. de Souza-Pinto

**Affiliations:** Department of Biochemistry, Chemistry Institute, University of São Paulo, São Paulo, São Paulo, 05508-000, Brazil; Institute of Mathematics and Statistics, University of São Paulo, São Paulo, São Paulo, 05508-000, Brazil

**Keywords:** aging, gene expression, co-expression network analysis

## Abstract

Transcriptomic changes occur with age, but the extent of their similarities across tissues is not clear. Previous studies have identified no similarity in age-modulated genes in different tissues. In this study, we sought to identify transcriptional changes with age across tissues using differential network analysis, with the premise that differential expression analysis alone is not capable of detecting all the changes in the transcriptional landscape that occur in tissues with age. Our results show major transcriptional alterations not detected by differential expression analysis that can be detected by differential connectivity analysis. Combining these two analyses, we detected genes changing across tissues, enriched in “RNA splicing” and “RNA processing”, and highly connected in protein-protein interaction networks. Co-expression module analyses demonstrated that other genes with tissue-specific variations with age are enriched in pathways that combat accumulation of aberrant RNAs and proteins, which are caused by defective splicing. Additionally, tissues displayed a major reorganization of their genes’ connectivities with age, with most demonstrating convergent connectivity patterns. Our analyses identify genes and processes which transcriptional changes are conserved across tissues, demonstrating a central role for RNA splicing and processing genes and highlighting the importance of differential network analysis for understanding the ageing transcriptome.

## INTRODUCTION

Aging is a multifactorial process that leads to alterations in several essential pathways, resulting in an increased risk of death. Nevertheless, despite its broad nature, the underlying mechanisms leading to the aging phenotypes are not well understood.

One pressing question in the field is whether tissues age similarly or differently. For instance, analysis of the methylation status of a small amount of CpG sites (epigenetic clocks) allows predicting an organism’s age with remarkable accuracy^1-5^. Notably, some of these epigenetic clocks are pan-tissue, developed with data generated from several tissues and representing a pattern of methylation changes common across several tissues^1; 3^. But despite the success in finding conserved DNA methylation patterns across tissues, finding such patterns in gene expression has proven to be more challenging. Several studies have compared age-related changes in gene expression across tissues in humans^6-10^ and rodents^6; 11^, with inconclusive results. While some report no overlap in differentially expressed genes with age between tissues, others find subtle commonalities between a few tissues at a process level, but not at the gene level. Therefore, the question of whether tissues age differently or if there are common aging alterations reflected in conserved tissue-independent transcriptomic changes remains.

Gene expression profiles can vary not only in their absolute expression values, but also in the genes’ co-expression patterns. Gene co-expression is typically represented as a correlation that measures how coordinated the expression of two genes are. Gene expression levels and gene co-expression patterns, however, do not have a direct relationship, and genes that vary in one aspect may not vary in the other. Given that previous works on gene expression and aging focused mostly on analyzing absolute gene expression changes, it is relevant to ask whether genes that do not change expression levels with age may have altered co-expression patterns. In principle, a gene does not need to change in its expression to have altered activity.

Weighted co-expression networks, constructed with pairwise gene expression correlations, provide a robust framework for analyzing system-level changes between different conditions^12^. In these networks, genes are represented as nodes, and vertices indicate pairwise relationships between genes. These relationships – essentially, gene expression correlations – provide a weighted measurement of the strength of the interaction between two genes. Gene co-expression networks have been successfully used in analyzing biological data, including in the aging context^13-16^.

Moreover, networks allow the calculation of different network structural measurements that can serve as metrics for comparing networks. One such metric is network connectivity, which represents the sum of the network adjacencies (vertices) for each gene. Therefore, connectivity in a gene co-expression network represents a measurement of each gene’s general correlation with all other genes in the network. In addition, eigenvector centrality scores nodes based on the centrality scores of the other nodes they are connected to, such that a node with a high eigenvector centrality is connected with nodes with high scores. Together, these network parameters reflect distinct biological information than expression levels.

In this work, we analyzed changes in gene expression and co-expression networks with age using RNA-Seq data from eight tissues from the Genotype-Tissue Expression (GTEx) project^17^. Our results demonstrate that changes in gene co-expression reveal alterations that cannot be revealed by analyzing changes in gene expression alone, and that when both analyses are combined, a more robust understanding of the regulation of gene expression during aging arises, showing conserved genes and processes, tied to specific protein complexes, being modulated with age across different tissues in humans.

## METHODS

### Dataset

Gene expression data (in TPM – transcripts per million) were obtained from the Genotype-Tissue Expression (GTEx) project (v8 release), which has *postmortem* RNA-seq data from 948 donors aged 20 - 70, from 54 tissues, totaling 17,382 samples. To allow the division of the dataset into age ranges with an adequate number of samples in each age range, we selected only tissues with at least 800 samples (blood, brain, adipose tissue, muscle, blood vessel, heart, skin, and esophagus). Tissues were analyzed at the level defined by the SMTS sample attribute in the GTEx annotation files.

### Differential expression analysis with linear mixed model

Differentially expressed genes were calculated for each tissue separately, using the following linear mixed model:

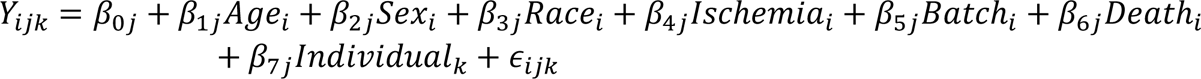

where, Y_ijk_ = expression level (log2 normalized TPM + 1) of gene *j* in sample *i* of individual *k*, *Age_i_* = age of sample *i*, *Sex_i_* = sex of sample i, *Race_i_* = race of sample *i*, *Ischemia_i_* = time of ischemia of sample *i*, *Batch_i_* = experimental batch in which sample *i*’s gene expression was measured, *Death_i_* = type of death, in the Hardy scale, of the donor from which sample *i* was collected, and *ε_ijk_* = error term (assumed to be normally and independently distributed with var(ε) = σ^2^I). Individuals are considered a block random effect, considering that some tissues have more than one sample for the same individual. Ischemic time was divided into 300-minute intervals to standardize the procedure with the removal of confounding factors, done in a later step. Random effects (Individual_k_) are assumed to be normally and independently distributed with var(*Individual*) = σ_ind_^2^I.

Genes with no or small variation in expression were filtered out of the analysis. Selected genes had expression higher than 0.1 TPM in at least 20% of the samples. To be considered differentially expressed with age, genes needed to satisfy an FDR^18^<0.1 or FDR<0.25, depending on the analysis, considering BH multiple test corrections. The model was fit using the nlme package in R^19^.

### Removal of confounding factor

For network analyses, expression was corrected by confounding factors using ComBat, from the sva R package^20^. Since ComBat can only correct for one effect at a time, effects were removed iteratively, following the order: sex, race, experimental batch, ischemic time, and death type. Given that ComBat does not accept continuous variables, ischemic time was divided into 300-minute intervals.

### Differential connectivity analysis

The dataset was divided into nine 6-year age ranges, except for the last one, with a 3-year range, to ensures that the smallest sub-dataset has at least 20 samples, the minimum recommended for WGCNA. Connectivity and eigenvector centrality for each gene were calculated in each age range using the *igraph* R package^21^. For comparison between networks, the connectivities in each age range were ranked, and a linear regression was applied to detect differences in ranked connectivities across the nine age ranges in the two metrics. Genes were considered differentially connected/eigenconnected with FDR<0.1 or FDR<0.25, depending on the analysis.

### Co-expression network construction and consensus module identification with weighted gene co-expression network analysis

Gene expression data for each tissue and age range, filtered for known confounding factors were used as inputs for the WGCNA package in R^22^. WGCNA constructs weighted gene co-expression networks using a soft threshold and fitting the data into a scale-free topology model. The chosen soft-thresholding powers for each tissue were: 6 (blood), 6 (brain), 9 (adipose tissue), 12 (muscle), 8 (blood vessel), 12 (heart), 6 (skin), and 6 (esophagus). We used the consensusBlockwiseModules function for module detection, with networkType set as “unsigned”. All parameters were kept at their default values. Sub-modules were defined by overlapping the positive and negative portions of the sets of genes of the three metrics (positive and negative DEGs, DCGs, and DECGs) with the modules defined by consensus modules analysis.

### Functional enrichment analysis

All enrichment analyses were carried out with the topGO R package^23^, using the classic Fisher algorithm. Annotations for each gene were retrieved from Ensemble 110^24^. Multiple test correction was performed using the BH method. Terms were considered enriched with FDR<0.1.

### Cross-tissue analysis of connectivity changes

Cross-tissue connectivity change analysis included only the 14,489 protein-coding genes that were present in the analyses of the 8 tissues (i.e. that were not filtered out the initial data filtering stage). In addition, we adjusted the connectivity ranking, since the connectivities were calculated and ranked including non-coding genes, preserving the effect of these genes in the connectivity. However, as non-coding genes represent around half of the genes available in the dataset, and their connectivities are, in general, small when compared to the protein-coding genes, they are heavily concentrated in the first half of the connectivity rankings, what caused large distortions in the histograms, making visualization difficult. Thus, we carried out a new ranking considering only the protein-coding genes used in this analysis.

For each tissue, reference histograms of the connectivity ranks of the DCGs of that tissue (called the reference tissue) were created. Given an x-axis representing all the connectivity ranks in bins of 100 ranks, the reference histograms show the distribution of the connectivity of all genes analyzed. For each tissue, four reference histograms were constructed: positive and negative DCGs in age range 1, positive and negative DCGs in age range 9.

With the reference histograms, histograms were constructed where for each gene considered a DCG in a reference tissue, the connectivity ranks of these genes in other tissues (called target tissues) are presented, regardless of whether they are DCGs in these tissues or not. In cases where they are DCGs in both the reference and the target tissue, the regions representing them in the histograms are colored (in red for positive DCGs and blue for negative DCGs).

### Comparison of enriched terms across sub-modules

To compare the results of the enrichment analyses between different sub-modules, we listed all enriched terms with FDR<0.1 in any submodule and construct a binary matrix with enriched terms in the rows and sub-modules in the columns. Entries had a value of 1 in the matrix if a term was enriched in the respective submodule. The terms were filtered: i) terms found in less than four tissues were removed, to focus on processes common across tissues; ii) terms found in the first four and last six levels of the Biological Process Gene Ontology tree were removed, with the rationale of removing terms that were too general or too specific to yield relevant biological information; and iii) we removed all the columns (sub-modules) that did not display any entries after the previous filtering. After filtering, 1,956 terms were included in the matrix. These steps ensure that: i) our matrix of terms per sub-module has a higher density of 1 values, making it easier to cluster, interpret and present; and ii) reduce the number of term, which is more suitable for the analysis of semantic similarity and presentation of semantic distances by Revigo^25^.

This binary matrix was clustered into 8 clusters with the *hclust* function from the stats package in R^26^, using the *Ward.D* clustering algorithm^27^. The results were presented as a heatmap using the *heatmap.2* function from the *gplots* package in R. The same list of enriched terms that make up the rows of the heatmap were summarized using the online tool Revigo^25^, which summarizes lists of Gene Ontology terms according to semantic similarities and creates a chart where the distances between points represent the semantic distances between terms. We also included an additional value representing the number of tissues in which each term is enriched (considering any sub-module in that tissue). Our list of terms was summarized using the following parameters in Revigo: *SimRel* semantic similarity algorithm, “Small” option for the size of the final list, “Yes” for removing obsolete terms, “*Homo sapiens*” for the species, and “Higher value is better” for the interpretation of the additional value (in our case, the number of tissues in which the enriched term is present). The R code for reproducing the chart was downloaded, and, within R, a variable containing an ID of the cluster in which that term was clustered by the *hclust* function was added. The final chart was plotted using the number of tissues as a variable to define the size of the points, and the cluster variable to define the coloring of the points.

### Protein-protein interaction network analysis with STRING

The PPI networks were analyzed with STRING^28^, using the STRING extension for Cytoscape^29^. For the construction of our networks, we used scores of at least 0.7, considered high in STRING.

### Gene overlaps

Gene overlaps and overlap significance statistics were calculated using the *GeneOverlap* R package^30^.

## RESULTS

### Detection of tissue-specific differentially expressed and connected genes

We used data from the GTEx project (v8), which has whole transcriptome data for up to 54 tissues from 948 *post-mortem* donors. To maximize statistical power, we aggregated sub-tissues and selected only tissues with at least 800 samples: blood, brain, adipose tissue, muscle, blood vessel, heart, skin, and esophagus. The data was processed as described in Methods. Figure S1A shows a general schematic of the data processing.

The differential expression and differential connectivity analyses were carried out as described. For the differential connectivity analyses, the dataset was divided into 9 age ranges (Figure S1B), and networks were constructed using WGCNA^22^. Network connectivity and eigenvector centrality were calculated using *igraph^21^*, resulting in nine connectivity measures per gene in each tissue across the nine age ranges. To compare the networks, we ranked the connectivities in each network and applied a linear regression on the nine age ranges to detect differences in ranked connectivities. All results obtained from the 3 regressions are found in Supplemental Data 1. Genes were considered altered in these metrics with FDR<0.1.

The number of differentially expressed genes (DEGs), differentially connected genes (DCGs), and differentially eigenconnected genes (DECGs) varied considerably by tissue (Figure 1A). Notably, many tissues demonstrated relative stability in either the differential expression analysis or one of the connectivity analyses. Blood, brain, muscle, and blood vessels all fit into this category, having many DEGs or DCGs/DECGs, but not both simultaneously. Adipose tissue had a high number of significant genes in all analyses, but a clear tendency towards having much more DEGs than DCGs. Only heart, skin and esophagus had relatively similar numbers of DEGs and DCGs/DECGs, though very few in heart.

**Figure 1.**
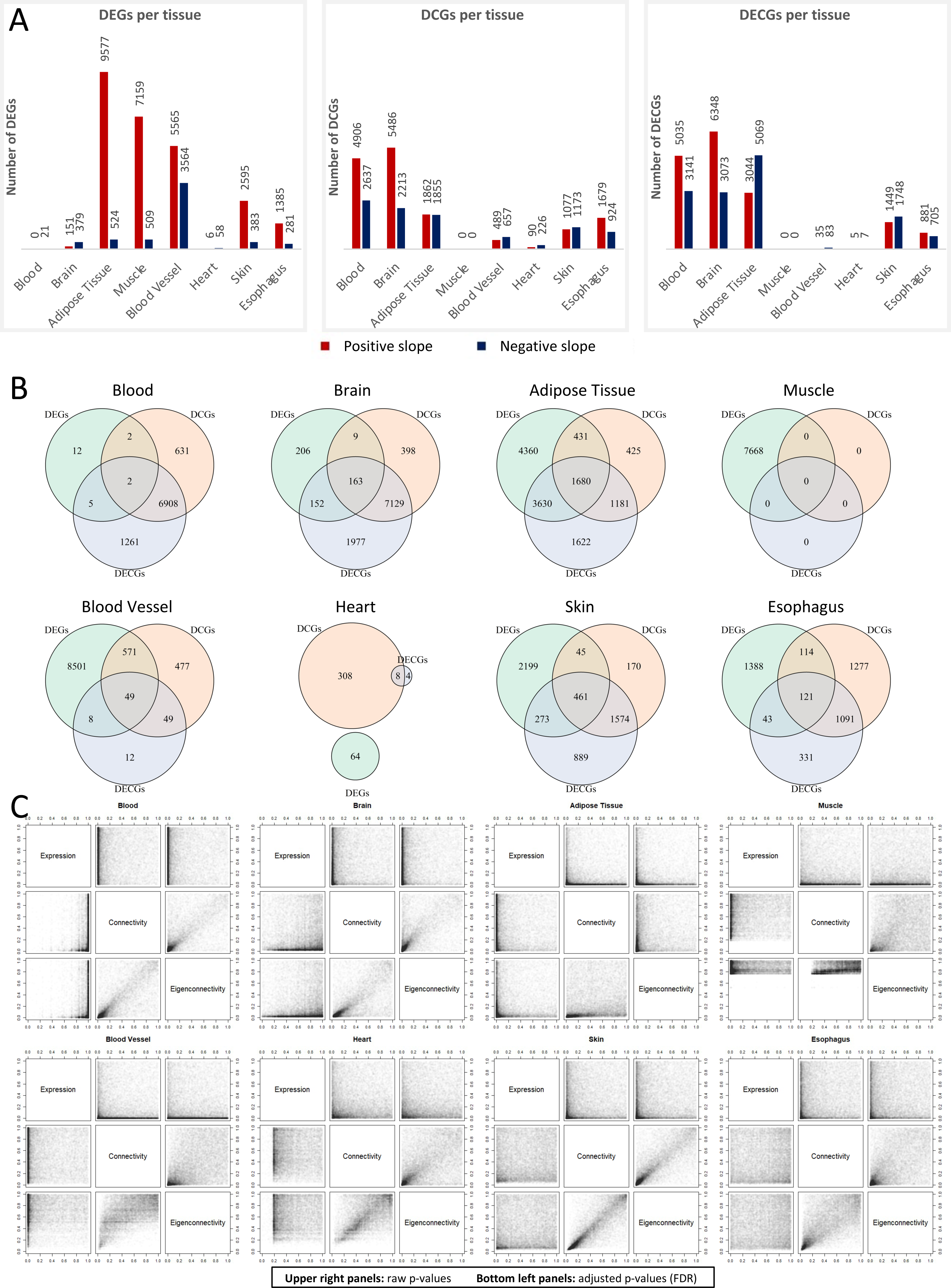
Altered genes per tissue. A) DEGs, DCGs, and DECGs per tissue. Red and blue bars represent genes with positive and negative slopes, respectively. All metrics considered FDR < 0.1. B) Overlaps between DEGs, DCGs, and DECGs in each tissue. C) Distribution of raw p-values and FDR-adjusted p-values in each tissue.

Only a small proportion of genes were simultaneously DEGs and DCGs/DECGs (Figure 1B). Adipose tissue and blood vessels had the highest overlap of genes detected in the three metrics. However, even in these tissues, the overlap between DEGs and DCGs/DECGs was less than half of the total genes detected. Looking at the p-values distribution of the three regressions against one another (Figure 1C), there is a greater concordance between the p-values of connectivity and eigenconnectivity regressions, but a great discordance between expression and any of the two connectivity metrics. These results show that even in tissues with a high number of significant genes in all metrics - such as adipose tissue, skin and esophagus – there is a clear tendency for a gene to change either expression or connectivity with age, but rarely both.

### Comparisons of altered genes across tissues

To compare whether the set of DEG or DCG/DECG is common across tissues, we counted, for each gene, in how many tissues it is changing in each metric. The results show that many genes are changing across tissues (Figure 2A – first table), but the intersecting gene sets get smaller as the tissue count increases, with few or no genes common for more than 6 tissues, depending on the metric.

**Figure 2.**
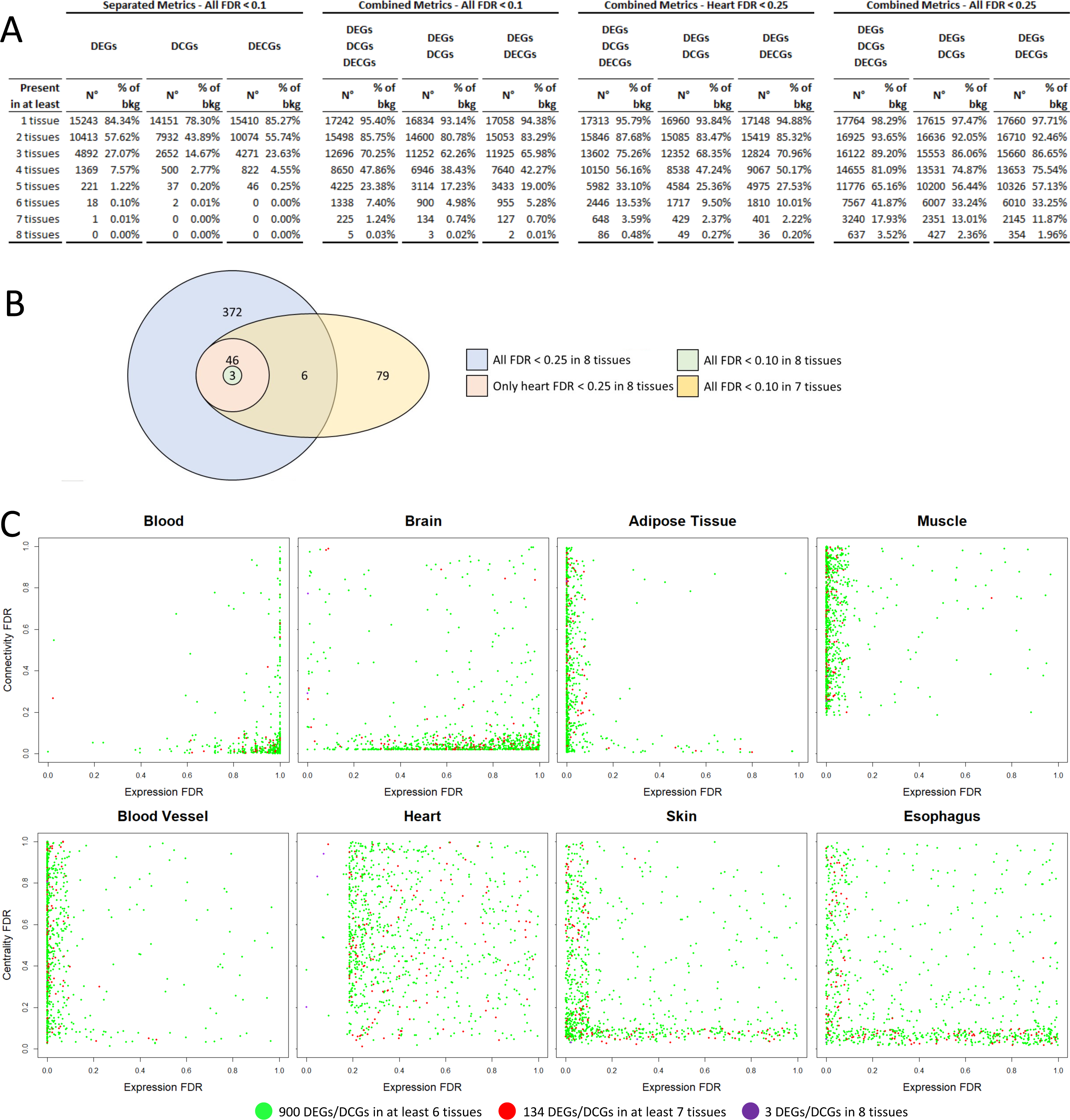
Intersect count. A) DEGs, DCGs, DECGs, and combinations of these metrics detected in sets of 1 up to 8 tissues. The tables display the number of genes in the set and their proportion in relation to the background of genes tested. B) Venn diagram of selected intersection sets. C) Scatterplots with FDR distributions in both regressions with intersecting genes highlighted. Green points represent the 900 DEGs/DCGs in at least 6 tissues, red points represent the 134 DEGs/DCGs in at least 7 tissues, and purple points represent the 3 DEGs/DCGs in 8 tissues.

However, given the discordant nature of the differential expression and differential connectivity regressions (Figure 1C), we asked whether genes that are DEGs in some tissues might be DCGs or DECGs in other tissues, with the rationale that both analyses detect changes in transcriptomic behavior, even if the type of change is different. For that, we considered three scenarios: whether i) the gene changed in any metric (DEG/DCG/DECG); ii) the gene is a DEG and/or a DCG (DEG/DCG); or iii) the gene is a DEG and/or a DECG (DEG/DECG).

Considering these scenarios, we identified 225 DEGs/DCGs/DECGs, 134 DEGs/DCGs, and 127 DEGs/DECGs common to at least 7 tissues (Figure 2A – second table), indicating that even though most genes changing with age are tissue-specific or shared between a few tissues, when intersections between tissues consider different metrics to assess transcriptomic behavior, a core of similar cross-tissue genes is uncovered.

The small intersection in 8 tissues is clearly skewed by heart. Interestingly, the distribution of raw p-values for heart displays a pattern not in line with the expected if the null hypothesis was true for all genes (none are changing) (Figure S2), as it can be observed for blood in the differential expression regression (Figure S2). It is clear, however, that the low p-values in the heart regressions were not as low as they were in other tissues. Since tens of thousands of tests were carried out, the multiple-test correction greatly penalizes the lowest p-values, which, in heart, are not as low as in other tissues. While a typical scenario of FDR correction with so many tests when there are close to no significant genes can be observed in the FDRs density histogram for blood in the differential expression regression, the same density histogram in heart displays a very different pattern, with most FDRs clustering around 0.20-0.25, which suggests a non-random distribution of p-values, raising the question of whether genes that cluster around 0.20-0.25 FDR in heart are meaningful.

Thresholds such as 0.05 and 0.1 are commonly used to define statistical significance. However, they are not based on theoretical justification, but on tradition and a necessity to define a standard in the absence of a theoretically defined threshold. While having such standard is necessary, sometimes it is essential to analyze results using broader statistical reasoning. Thus, we hypothesized that if the heart indeed has a similar core of genes intersecting with other tissues, then they must be among the lowest ranked FDRs in heart, probably in the 0.20-0.25 interval, where many genes are clustered (Figure 1C – Heart, bottom left panels). Even if these genes fail significance using a traditional threshold, if the lowest ranked FDRs in heart happen to be the very same that are also changing in other tissues, this strengthens to the idea that there is a core of genes that changes across all tissues.

To explore this, we analyzed two additional scenarios for the intersection analysis: considering genes significant at FDR<0.25 only in heart, or FDR<0.25 for all tissues.

The last scenario considers FDR<0.25 for all tissues, but since we are interested only in genes intersecting across all tissues, increasing the threshold for all and focusing only on the intersecting sets in 8 tissues may still lead to interesting results worth exploring, since if a similar set of genes is among the ones with the lowest p-values in all tissues, this is very unlikely to be a coincidence.

By increasing the threshold only for heart, we uncovered common genes with other tissues (Figure 2A – third table). By increasing the threshold for all tissues, we find a larger number of shared genes intersecting in 8 tissues (Figure 2A – fourth table).

Since the results for the connectivity and eigenconnectivity analyses were well correlated, further analyses focused on DEGs and DCGs. The sets of genes presented in Figure 2A are referred to as “intersection sets” hereinafter, and the specific sets of interest (with the combination of DEGs and DCGs) will be called: 3-set, 49-set, 427-set and 134-set, in reference to the number of genes in each set. Figure 2B shows a Venn diagram overlapping these sets of interest.

To further show this similarity in heart, we plotted the FDR distribution of genes belonging to the three sets encompassing more tissues using FDR<0.1 (900 DEGs/DCGs in 6 tissues, 134 DEGs/DCGs in 7 tissues, and 3 DEGs/DCGs in 8 tissues). Naturally, they cluster very strongly in the lower FDR extremities of the charts of the 7 tissues other than heart. However, as the results in Figure 2A indicate, they cluster in the lower FDR ranges in heart, even if at FDR thresholds higher than 0.1. This visual demonstration reinforces that those genes found in common among 6-7 tissues at FDR<0.1 are also among the lowest ranked p-values in heart.

### GO enrichment analysis of intersecting gene sets

We performed a GO enrichment analysis of the intersection sets using the *topGO* R package. Given that we are not interested in the tissue-specific changes, but in the common genes across tissues, we limit the results to the intersection sets comprised of less than 5000 genes (sets with less than 5000 thousand genes in the tables in Figure 2A). Enrichment results for all these sets are found in Supplemental Data 2, 3 and 4. Figure 3 presents the results for selected intersection sets. The sets shown are those with most tissues for which the enrichment results found significant terms in the Biological Process ontology, with a reasonable number of significant genes annotated to the term (see Methods for a more detailed description of the rationale for the choice of these sets). When the number of terms was too large for proper display in the chart, we choose to display the Biological Process terms and, if still necessary, remove the terms with fewer genes annotated to them.

**Figure 3.**
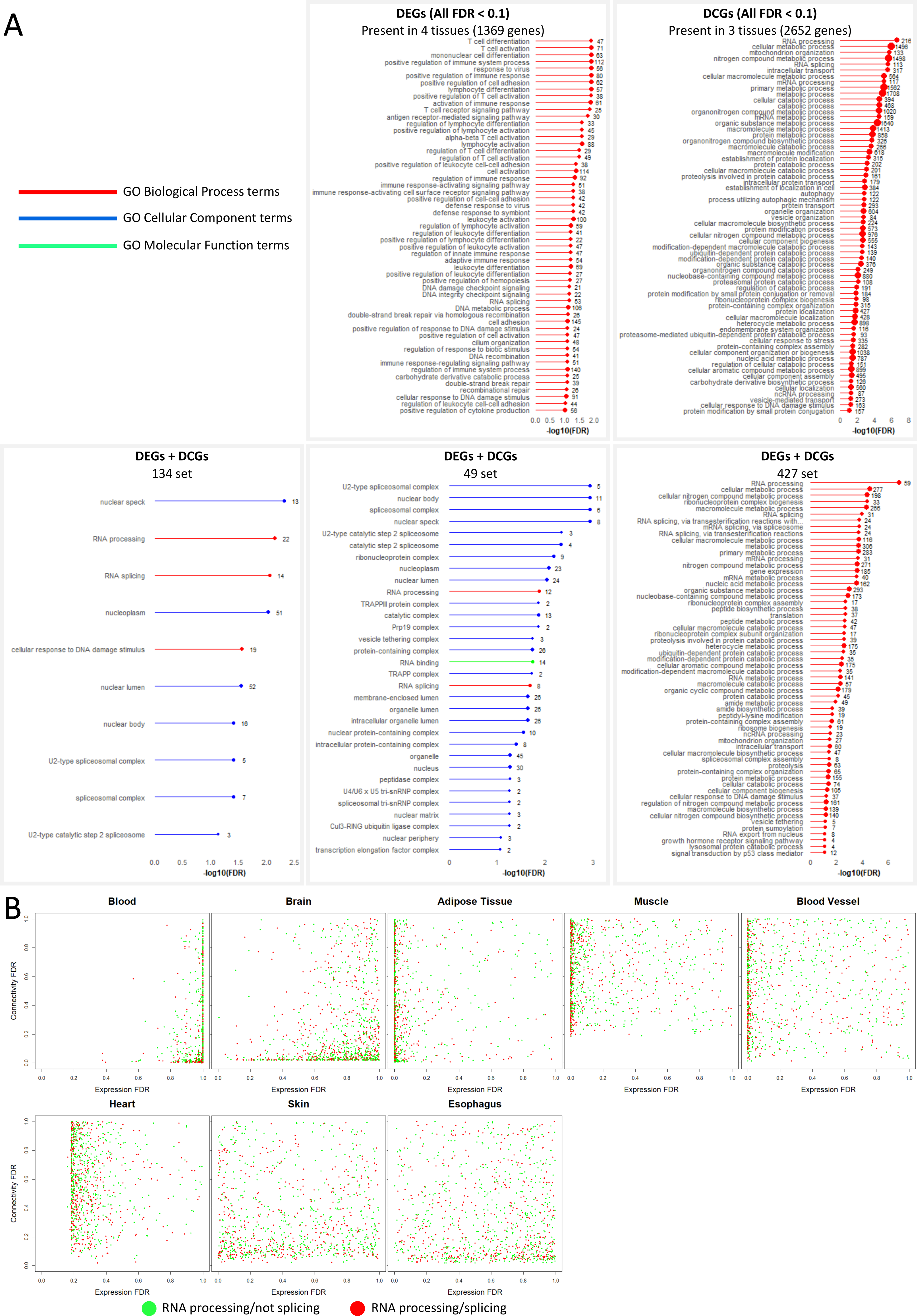
GO enrichment analysis of intersection sets. A) Enrichment of selected intersection sets. Terms significant at FDR < 0.1. In cases where too many terms were enriched, only Biological Process terms are shown. When that still yielded too many results, further filtering was done by removing the terms with the least number of genes. B) Distribution of genes annotated as “RNA processing” or “RNA splicing” in the FDR distribution charts.

While the DEGs chart displays immune system-related terms not found in the DCGs chart, they both contain similar terms related to RNA splicing and DNA damage response. Furthermore, while the set of DCGs in 4 tissues was not chosen because of the previously mentioned criteria, its enrichment analysis does show enrichment in Cellular Component terms for the spliceosomal complex (Supplemental Data 2), indicating that four tissues have good evidence for the enrichment of intersecting DEGs and DCGs in terms related to RNA splicing and DNA damage response.

The selected intersection sets in 7 or 8 tissues also show enrichment in RNA splicing, RNA processing and DNA damage response. The FDR distribution of all RNA splicing and processing genes corroborates the tendency of these genes to be in the areas of most significance in the charts of all tissues, and reinforces the tissue-specific discordant behavior of the genes in the two regressions. It is important to note that the term RNA splicing is a direct child of the term RNA processing in the Biological Process ontology tree, meaning that RNA splicing genes are contained in the RNA processing genes. DNA damage response is not found in the 49-set, but it is in the 427-set, along with other terms related to protein catabolism, autophagy, and ribosome biogenesis. Figure S3 presents the induced GO subgraph of the top 20 enriched terms in the 134-set.

In most tissues, genes in these sets are mostly upregulated in at least one of these two metrics (Figure S4), except for blood vessels, with a mixed profile, and heart, in which most genes are negative DEGs. Furthermore, negative DEGs, including in the 427-set, tend to be highly expressed in younger samples (Figure S5). In fact, in blood vessels and in heart, tissues with a higher proportion of negative DEGs, both overall and in the intersection sets, most of the negative DEGs remain highly expressed in older samples, suggesting that most negative DEGs remain highly necessary.

Connectivity changes between young and old samples tended to be more dramatic, with DCGs in the 427-set and in the splicing genes set tending to become more highly connected at older ages. The most obvious exception seems to be skin, where most splicing and 427-set genes decrease their connectivity (Figure S5). The 134-set and the 49-set show a similar pattern (Figure S6).

### Cross-tissue analysis of connectivity changes

The fact that some tissues have a tendency towards more DEGs or DCGs with age is puzzling. Even in skin and esophagus, which had a more balanced number of DEGs and DCGs, the genes changing in one metric are mostly not changing in the other. In order to explore this phenomenon, we analyzed the distribution of connectivity ranks of DCGs in each tissue against the connectivity ranks of the same genes in other tissues (see Methods).

The results indicate that positive DCGs in each tissue tend to be among the lowest or middle ranked genes in their respective tissues in younger samples (Figure 4A). However, when we compare the connectivity rank histogram of these same genes in other tissues, we find they tend to be much more highly ranked in the other tissues. For example, positive DCGs in blood tend to be among the low and middle ranks in terms of connectivity. However, the same genes that are positive DCGs in blood tend to be more highly ranked in other tissues. Brain, adipose tissue, muscle, blood vessels, and heart, present a significant skew to the right on the histograms. While less obvious, skin and esophagus also show higher concentration of genes among higher ranks. Negative DCGs present similar behavior, with an inverse logic: negative DCGs in a tissue tend to be already less ranked in other tissue in which they are not negative DCGs. The only exception being skin, since its negative DCGs seem to be highly ranked in many other tissues.

**Figure 4.**
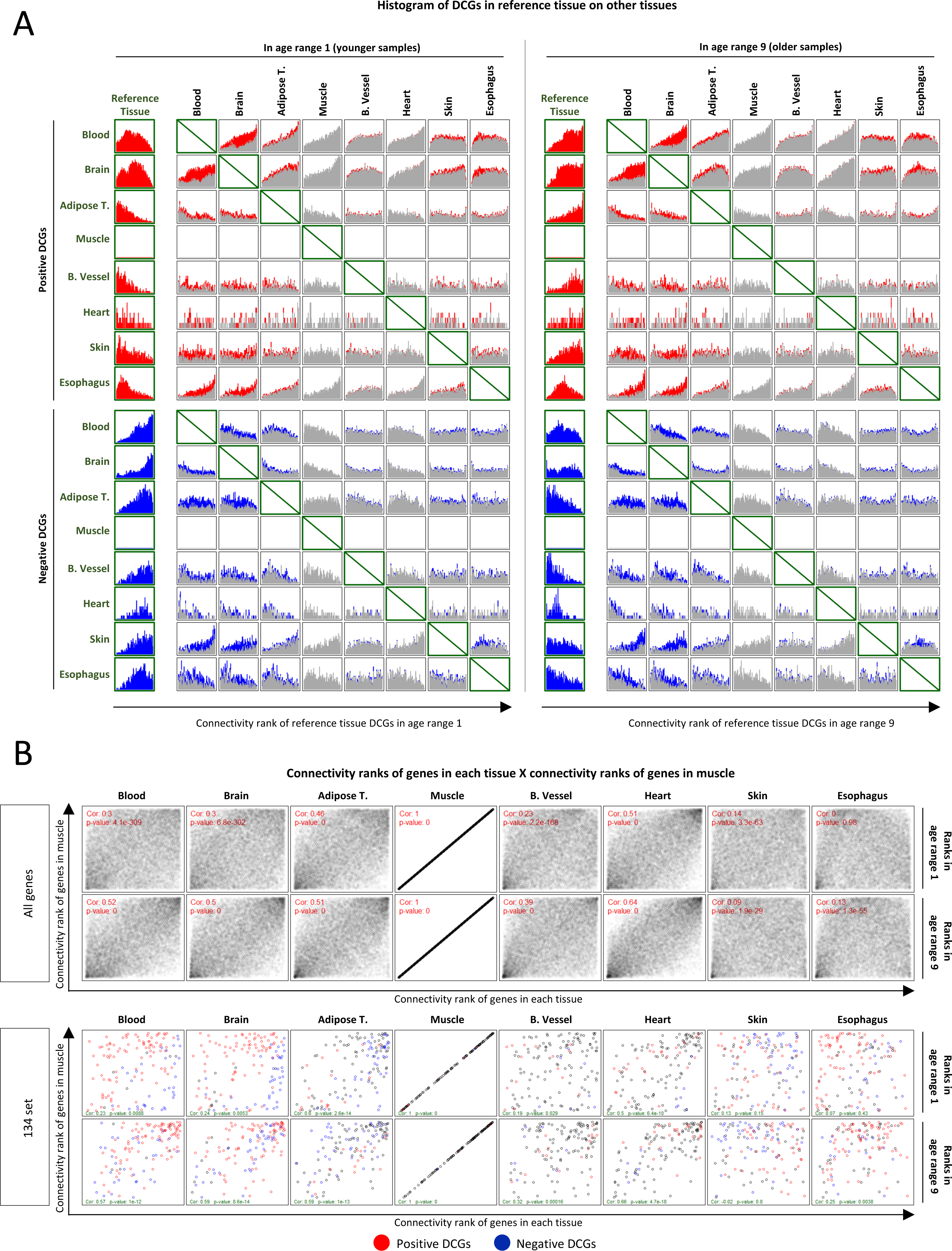
Cross-tissue analysis of DCGs. A) DCGs in reference tissues projected on other tissues in younger and older samples. Colored portions of the histogram represent genes that are positive (red) or negative (blue) DCGs both in the reference and in the target tissue. Grey portions represent genes that are DCGs in the reference tissues but not in the target tissue. B) Connectivity ranks of genes in each tissue X connectivity rank of genes in muscle. Muscle was chosen as a reference because it does not have any DCG at FDR < 0.1.

In older samples, the DCGs change their rankings as expected, such that the connectivity ranking pattern of these genes closely resembles the pattern that they already displayed in other tissues, suggesting that the gene connectivity is getting reorganized in some tissues, and that in tissues where these genes are not changing their connectivity, it might be because connectivity was already roughly at the biological “ideal” level.

This apparent convergence of connectivity ranks is clear when plotting the connectivity ranks of all genes in each tissue against the connectivity ranks of the same genes in muscle (Figure 4B). Muscle was chosen as a reference because it lacks DCGs. The correlations between the connectivity ranks of genes in any tissue compared to muscle tend to increase considerably with age in most tissues. Only skin displayed a slight decrease in correlation. Similar results can be seen when plotting only the 134-set (Figure 4B), the-49 set, the 427-set, or all RNA splicing and RNA processing genes (Figure S7).

No clear convergence can be observed using DEGs instead of DCGs (Figures S8 and S9). The only noteworthy observation is that the correlation between the expression of genes across tissues is high (0.5-0.65). When we consider only the intersection sets or just RNA processing/splicing genes, the correlation is even higher (>0.8 in many cases), suggesting that the intersection sets uncovered here and the broader set of all RNA processing/ RNA splicing genes behave more similarly across tissues than the universe of all genes, both in younger and in older samples, despite some being DEGs. This contrasts with the results comparing DCGs across tissues, which show remarkable levels of reorganization in some tissues and a converging profile in all tissues but skin.

### Consensus network and GO enrichment analysis of age-related sub-modules

While only a few genes seem are common across 7 or 8 tissues, we asked whether genes common in fewer tissues could belong to similar biological processes. For that, we used WGCNA to build an unsigned consensus network for the nine age ranges in each tissue, which allowed identification of conserved gene co-expression modules that could be found in all nine datasets. After obtaining these modules, they were overlapped DEGs, DCGs and DECGs, and divided into positive and negative portions. This led to the definition of sub-modules, where each sub-module is composed of the positive or negative DEGs, DCGs, or DECGs that are contained in that module. By dividing the DEGs/DCGs/DECGs using the modular structure provided by WCGNA, we can group them into sets that are likely to be related to the same biological processes and, therefore, identified based on a functional enrichment analysis.

Since the sub-modules enrichment generated many lists of enriched terms, we summarized them first in a heatmap format (Figure 5A and S10) and used Revigo to display the results (Figure 5B) (see Methods for a detailed description). Complete enrichment results per module are found in Supplemental Data 5, 6, and 7.

**Figure 5.**
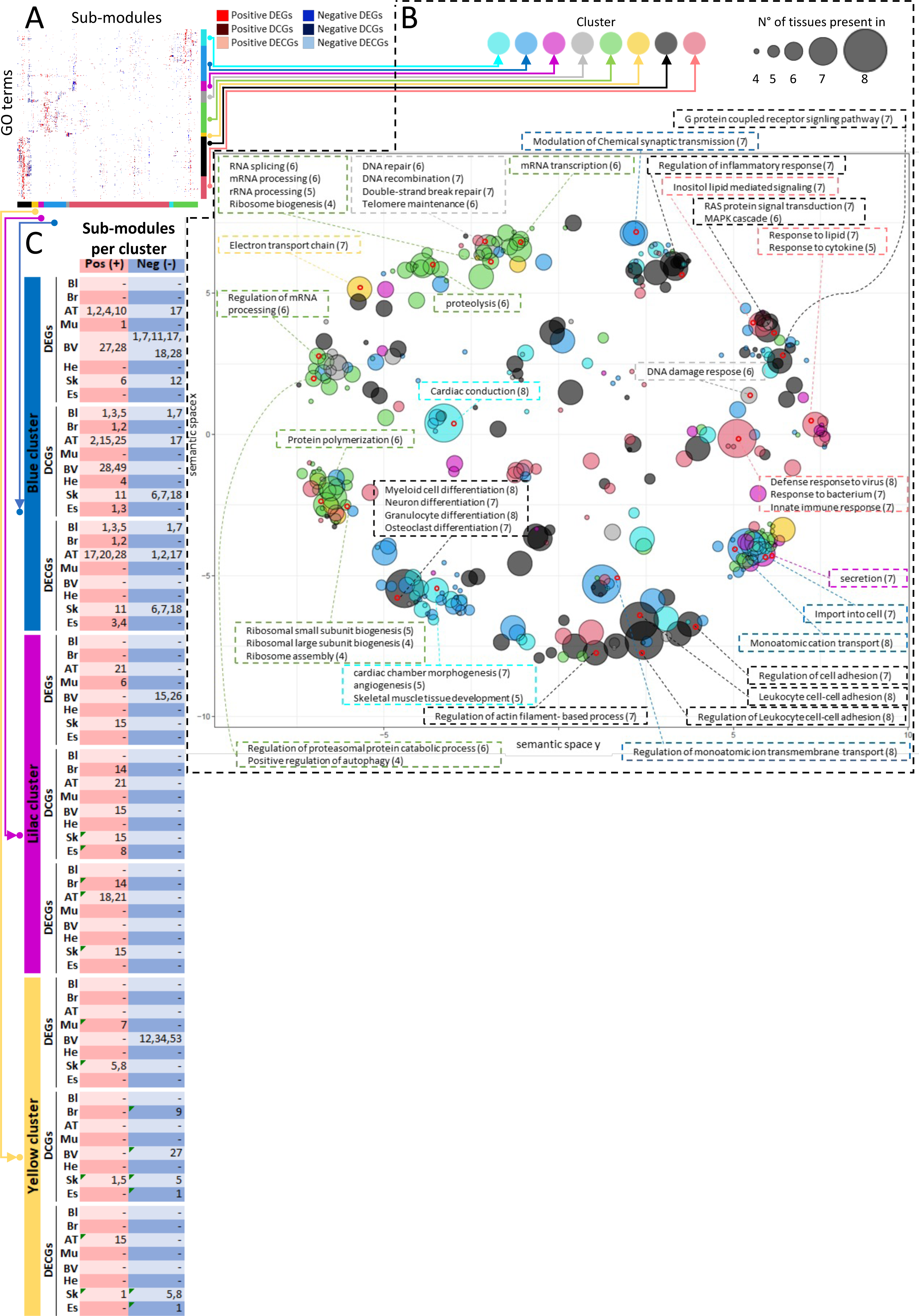
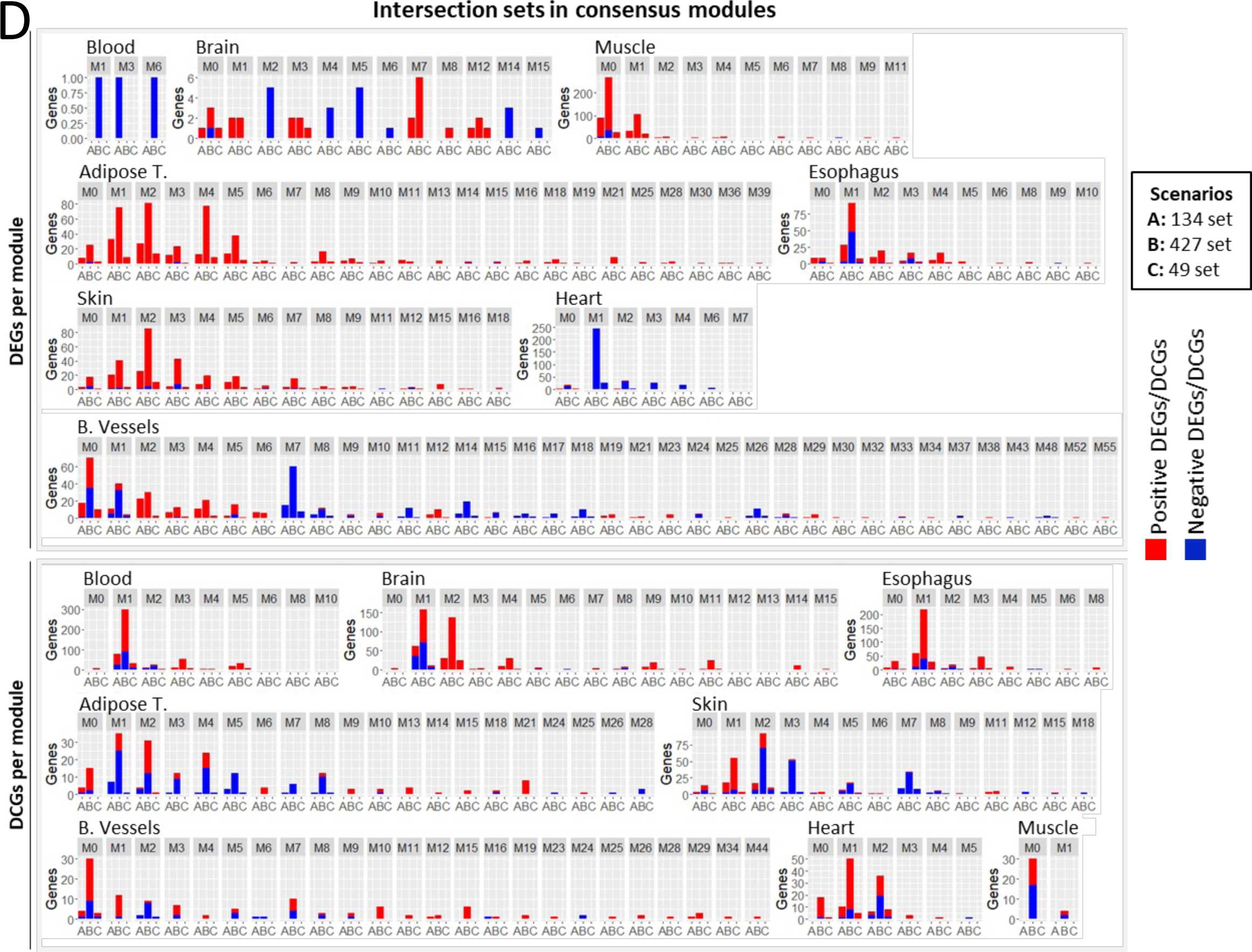
Consensus modules analysis. A) Unlabeled heatmap of similarity between terms enriched in sub-modules. B) Semantic similarity of the terms in the heatmap with Revigo. The size of the dots on the chart represents the number of tissues in which that term is found, and the coloring of the dot represents the cluster of terms/rows in which that term is found. Both the size and coloring are based on the term chosen as representative of the cluster. Terms contained in some points in different regions of the chart are highlighted and described in the text boxes. The numbers in brackets next to the terms represent the number of tissues in which that term is enriched (which may or may not be equal to the value for the representative term of the same point). C) Sub-modules enriched in terms contained in the green and gray terms/rows clusters. D) Overlap between consensus modules and genes in the intersection sets. Red and blue portions of the bars represent, respectively, positive and negative DEGs/DCGs.

The green rows/terms cluster contains sub-modules enriched in RNA splicing and processing, the same processes enriched in the intersection sets (Figure 5A and S10). Inspecting the heatmap and Revigo results (as well as Figure S10 and Supplemental Data 5, 6 and 7), we find that, in addition to the RNA splicing and processing terms, other processes were frequently enriched in the sub-modules in the green cluster, such as proteolysis, protein catabolism, autophagy, and ribosomal biogenesis. Additionally, the grey cluster show a large sub-module overlap with the green cluster, and contains terms related to DNA repair and DNA damage response, suggesting that these processes are changing in a coordinated fashion during aging, already indicated by the enrichment of the 427-set (Figure 3).

Figure 5C shows the sub-modules that concentrate terms contained in the green and grey rows/terms clusters. These sub-modules seem to group in the blue, lilac, and yellow columns/sub-modules clusters. In the table, we present the sub-modules numbering, and identify if they are composed of DEGs, DCGs or DECGs; positive or negative.

Subsequently, we counted the number of DEGs and DCGs in the intersection sets in each module in each tissue (Figure 5D). Most genes in the intersection sets are grouped in the first (and largest) modules, which contain the sub-modules shown in the table in 5C (contained in the blue, lilac and yellow clusters of columns/sub-modules), which are enriched with terms related to RNA processing and splicing, proteolysis, autophagy and ribosomal biogenesis (green cluster of rows/terms); and DNA repair and DNA damage response (gray cluster of rows/terms).

On the other hand, heart and skin behave differently. We reasoned that this is to be expected for heart, since the DEGs/DCGs/DECGs chosen for the sub-modules were only those with FDR<0.1, which were very few. For skin, only M7 sub-modules are found in the blue cluster in Figure 5C. M1 and M5 skin sub-modules are present in Figure 5C, but are contained in the yellow sub-module cluster, which although it has a pattern of terms like the blue sub-module cluster, has a higher concentration of terms from the gray cluster, more related to DNA repair and DNA damage response. Although M2 sub-modules were not in the clusters in Figure 5C, by inspecting the enrichment results of the M2 sub-modules (Supplemental Data 5, 6 and 7), we found that no term was enriched in almost all the sub-modules of this module, explaining why they were not clustered in Figure 5A. The M3 module also has enriched terms in the sub-modules for positive DEGs, negative DCGs and negative DECGs (Supplemental Data 5, 6 and 7), although they are more enriched in terms related to ribosomal biogenesis, regulation of translation and RNA processing. Only in the sub-module of negative DECGs terms related to RNA splicing appear enriched at FDR<0.1.

These results show that although the genes in the intersection sets were the only ones changing across tissues, they are associated with modules that contain other genes that are changing in a tissue-specific manner, related to similar processes and which act in a coordinated fashion with the genes in the intersection sets.

### Protein-protein interaction prediction network analysis

Co-expression relationships do not identify the molecular mechanisms through which interactions manifest. Therefore, in order to determine whether there are physical associations between the protein products of these genes, we constructed protein-protein interaction (PPI) networks using STRING. The STRING database has not only curated protein-protein interactions but also has predictions of protein-protein interactions based on several methods. To analyze the interactions between the genes we found, we queried all of them in STRING, including the 427 genes found in the larger set, with the addition of the genes in the 134-set that were not already included in the 427-set. This added up to a total of 506 genes, of which 505 were identified in the STRING database. We constructed two networks: one displaying only the physical subnetwork (which links only proteins that share a physical complex) and another displaying the full STRING network, using a confidence score of 0.7 for the edges for both.

Results are presented in Figure 6. Nodes with no connections (singletons) were left out of the network. Physical complexes are highlighted by dashed lines around them, and nodes are colored according to which gene set they belong to.

**Figure 6.**
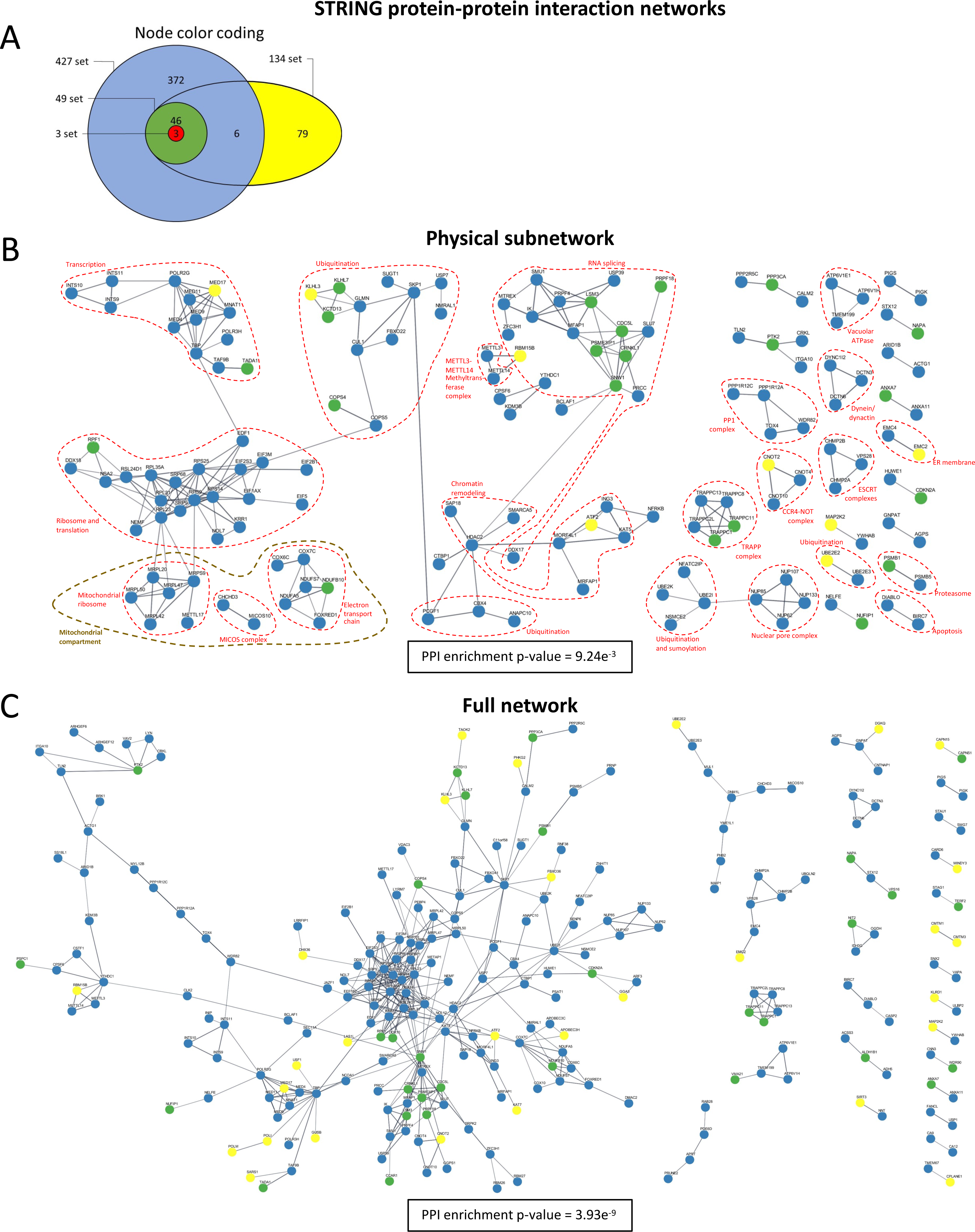
STRING protein-protein interaction networks. A) Color coding for the nodes in the networks indicating to which intersection set the corresponding genes belong. B) Physical interaction subnetwork. C) Full interaction network. Both networks include only links with a confidence score of at least 0.7 in STRING.

Several proteins found in the analysis have good evidence of being connected in physical complexes (Fig 6B). In fact, many of them cluster into very well-defined physical complexes, including several that interact in the spliceosome, as expected from our previous results.

Additionally, many ribosomal proteins (both cytosolic and mitochondrial) are present, along with translation initiation and elongation factors. Several proteins involved in transcription and chromatin remodeling are also clustered, as well as ubiquitination and proteolysis proteins. Most complexes are composed of proteins added with the flexibilization of the FDR threshold to 0.25 in all tissues. However, splicing-related proteins showed good physical clustering even using the 49-set and 134-set. Many additional proteins related to these processes, including RNA splicing, were left unconnected (as singletons) in the network due to not reaching the required confidence score in STRING or having more indirect connections. The PP1 complex highlighted is also involved in regulating RNA splicing^31^.

The complexes shown seem to delineate a continuous path, with the transcription of mRNAs being followed by RNA splicing and processing, and subsequent translation in the ribosomes, after which proteins are sent to the ER; during this process, mRNAs and proteins are being degraded. The degradation of proteins is represented in the network by proteins that promote the ubiquitination and tagging for degradation, as well as some protease-related proteins. In the mitochondria, in addition to the mitochondrial ribosome, there are also components of the electron transport chain. Interestingly, only components of Complexes I (NADH dehydrogenase) and III (cytochrome c oxidase) seem to be altered.

The full network (Fig 6C) displays the complexes seen in B, but with more proteins connected, with the larger cluster in the middle representing ribosomal proteins, and the cluster below RNA splicing factors.

## DISCUSSION

Several studies have tried to look for similarities in gene expression across tissues^6-11; 32-34^; with some employing WGCNA^13; 16^, but no similarities were found at the gene level, and very few were found at the process level.

Our work’s initial premise was that differential expression analyses are not capable of detecting all alterations that occur in the transcriptome with aging, and that a lot of information is lost when we only analyze changes in absolute expression levels. This lost information may be able to explain why there has been so much difficulty in finding clear common patterns in transcriptomic changes with aging across tissues, especially in terms of individual genes. We hypothesized that part of this information may be contained in changes in co-expression (reflected in network topological metrics). We did this under the hypothesis that some genes may not display differences in absolute gene expression levels with age, but may display differences in how they co-express with others. This yielded very different results from a traditional differential expression analysis, revealing patterns of transcriptomic changes that cannot be seen by looking only at absolute gene expression values, as some tissues with very few DEGs displayed a considerable amount of DCGs and DECGs, and vice versa.

Our results demonstrate that there is a core of genes enriched in RNA splicing and processing terms that are being altered across tissues, and that are well connected in PPI networks, but that these changes are either captured by differential expression or differential connectivity analyses, but rarely in both. This result is true for 7 tissues with FDR < 0.10, and for all 8 tissues with FDR<0.25.

This demonstrates that looking only at differential expression may lead to very incomplete results in some tissues. For instance, given the preponderance of DCGs in blood and brain, it is no surprise that no common set of DEGs would be found between them and other tissues. Any analysis comparing only DEGs using these two tissues would miss the reality that these tissues display transcriptomic changes with aging that are reflected much more heavily in co-expression/connectivity related changes than in absolute expression changes. Therefore, ignoring the differences in connectivity in some tissues may lead to a false notion that they do not display age-related transcriptomic changes in a common set of genes. When this logic is considered, the question ceases to be whether there is a common set of genes related to specific enriched processes changing across tissues with aging, and a new question arises asking why are these different tissues demonstrating transcriptomic changes in different ways.

At least in terms of DCGs, it seems that differences can be mostly explained by the fact that the connectivity of genes in each tissue seem to be reordering in such a way that their rankings are converging with aging. With skin being the only exception. This behavior could be an indication that a common stimulus is afflicting them. If the burden of defective spliced RNAs and consequent malformed proteins increases with aging, it seems reasonable that all tissues will have to reorganize themselves in order to deal with this same problem. However, given that different tissues will have different starting points regarding the connectivity of each gene, it makes sense that how many genes will change in a given tissue and the trajectory of these changes will be different according to tissue, even if the endpoint might be approximately the same. Some tissues might already have a specific configuration of connectivity of their genes while young that allow them to deal with higher burdens of aberrant splicing and malformed proteins. In contrast, others may be more susceptible to this type of alteration, and may need to promote a greater reorganization in the way their molecular network is structured in order to deal with this increasing burden. Therefore, a common stimulus may be conditioning different tissues to converge to a similar adaptive state.

One possibility for the exception in the skin might be that it is the only tissue analyzed that is exposed directly to the external environment, which might make the burden of exogenous factors much larger than in other tissues, creating a different pattern of reorganization. Nonetheless, there is mostly upregulation of DEGs in the intersection sets for skin, indicating that the burden of defective spliced RNAs may indeed be greater. The fact that many negative DCGs in skin are also positive DEGs (Figure S4), including RNA processing/splicing genes, indicates that they do require an increased activity with aging. Furthermore, there is a significant subset of genes in the intersection sets and RNA splicing sets that are positive DCGs, indicating, to some extent, a greater necessity for these genes.

Indeed, aging seems to be accompanied by a high incidence of aberrant splicing and intron retentions in *C. elegans*, fruit flies, mice, and humans^35-39^. Notably, Mariotti et al. (2022)^39^ showed an increase in these events in the GTEx dataset for several tissues. Most tissues they analyzed showed an increase in intron retention with aging. This includes heart and blood vessels, for which we detected decreased expression in the intersection sets. One possible explanation for the observation that these tissues do seem to have a higher aberrant splicing burden but less expression of splicing genes may be that the changes in connectivity ranks are taking the lead in adapting these tissues to this alteration. Since both tissues display a convergent pattern of connectivity ranks, these changes may be already boosting the adaptation to aberrant splicing as much as the tissue can. Furthermore, the negative DEGs in the intersection sets do not seem to result in a dramatic decrease in expression.

It is not possible, however, to know with certainty what causes this increased incidence of aberrant splicing. One possible hint is the presence of genes associated with DNA repair and DNA damage response. The idea that DNA alterations cause aging is one of the most classic theories of aging^40; 41^, and DNA damage is a hallmark of aging^42^. It is possible, therefore, that damage to DNA is leading to aberrant splicing. Indeed, there seems to be a connection between RNA splicing and DNA damage response^43^. Furthermore, previous studies have shown that higher expression of DNA repair genes is associated with species’ maximum lifespans^44-46^, and the transcriptional signatures associated with maximum lifespan seem to present similarities with those seem during aging in a tissue-specific manner^47^.

The processes clustering with RNA processing/splicing in the same modules are interesting because they all act in a coordinated fashion during the production cycle of new proteins and can be very easily analyzed concerning what we know about aging and the lifespan extension promoted by mTORC1 inhibition. Inhibition of mTORC1, through genetic or pharmacological means, is one of the most studied methods of life span extension and works in several organisms, including mice^48-63^. mTORC1 is a central regulator of autophagy^64^. Therefore, higher expression and relative connectivity of autophagy and proteolysis/protein catabolism genes, as indicated in our enrichment results, demonstrates that these mTORC1-related processes are more prominent during aging (see Figure 5 and Supplemental Data 5, 6 and 7). Loss of proteostasis is a hallmark of aging^42^, and aberrant splicing is a cause of malformed proteins, which demand more of the protein degradation mechanisms to try to remove these malformed proteins.

Lastly, ribosomal biogenesis has also been associated with aging^65^. The presence of these processes in the same modules suggests that they work in a coordinated fashion. Higher rates of splicing abnormalities may lead to more malformed proteins, which will demand more of the protein degradation machinery. Such aberrant mRNAs may create difficulties for the ribosomal machinery since they may take longer to translate completely, given the difficulties in folding. Therefore, we could speculate that this increased occupancy of ribosomes would end up requiring the biogenesis of more ribosomes, in order to decrease the amount of mRNA in line to be translated.

From the results described here, it is reasonable to propose a scenario in which the age-associated increase in aberrant mRNAs, proteins, and eventually organelles, which were negatively affected by malfunctioning proteins, may impose significantly on catabolic processes such as RNA catabolism, protein catabolism, and autophagy. Lifespan extension by mTOR inhibition may thus be working by inducing the clearance of these defective components. Since these clearance mechanisms seem to be upregulated with age, mTOR inhibition is boosting a naturally occurring attempt at adaptation by the cells.

## CREDIT AUTHOR STATEMENT

CMPFB: conceptualization, methodology, software, formal analysis, investigation, writing. AF: methodology, validation. NC S-P: conceptualization, writing, supervision, funding acquisition.

## ACKNOWLEDGEMENTS

This work was supported by Fundação de Amparo à Pesquisa do Estado de São Paulo (FAPESP) grant # 2017/04372-0 to NCS-P. CMPFB was supported by Conselho Nacional de Desenvolvimento Científico e Tecnológico (CNPq) grant # 140207/2018-0.

## DATA AVAILABILITY

The raw data used in the analyses reported in this study are available through the Genotype-Tissue Expression (GTEx) project at gtexportal.org/home.

---

**Figure S1.**
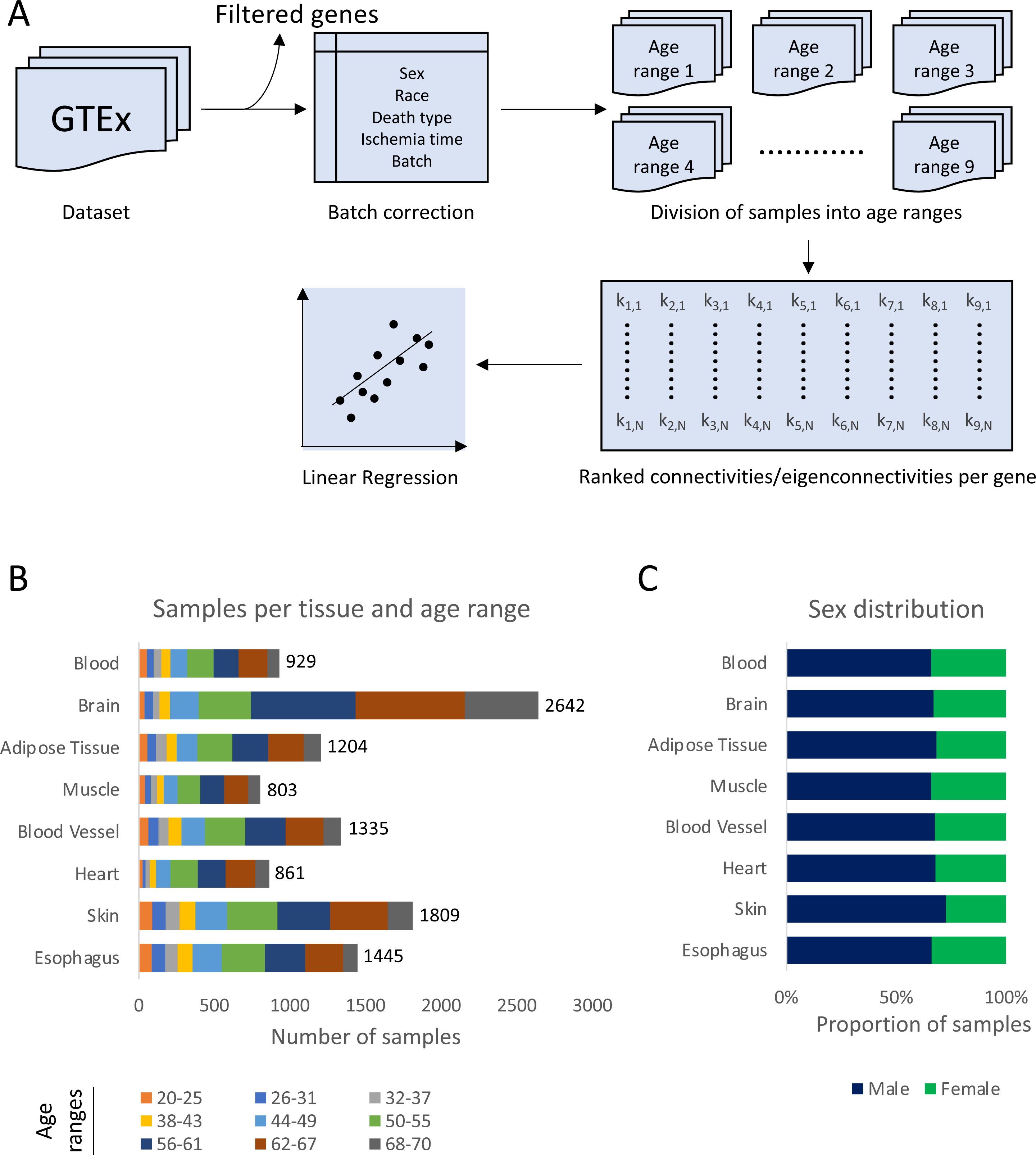
Dataset description. A) Schematics of the partition of the dataset into age ranges for the calculation of differential connectivity and eigenconnectivity. B) Number of samples per tissue per age range; age ranges account for 6 years each, except for the last one, which accounts for 3 years. C) Proportion of male and female samples in each tissue.

**Figure S2.**
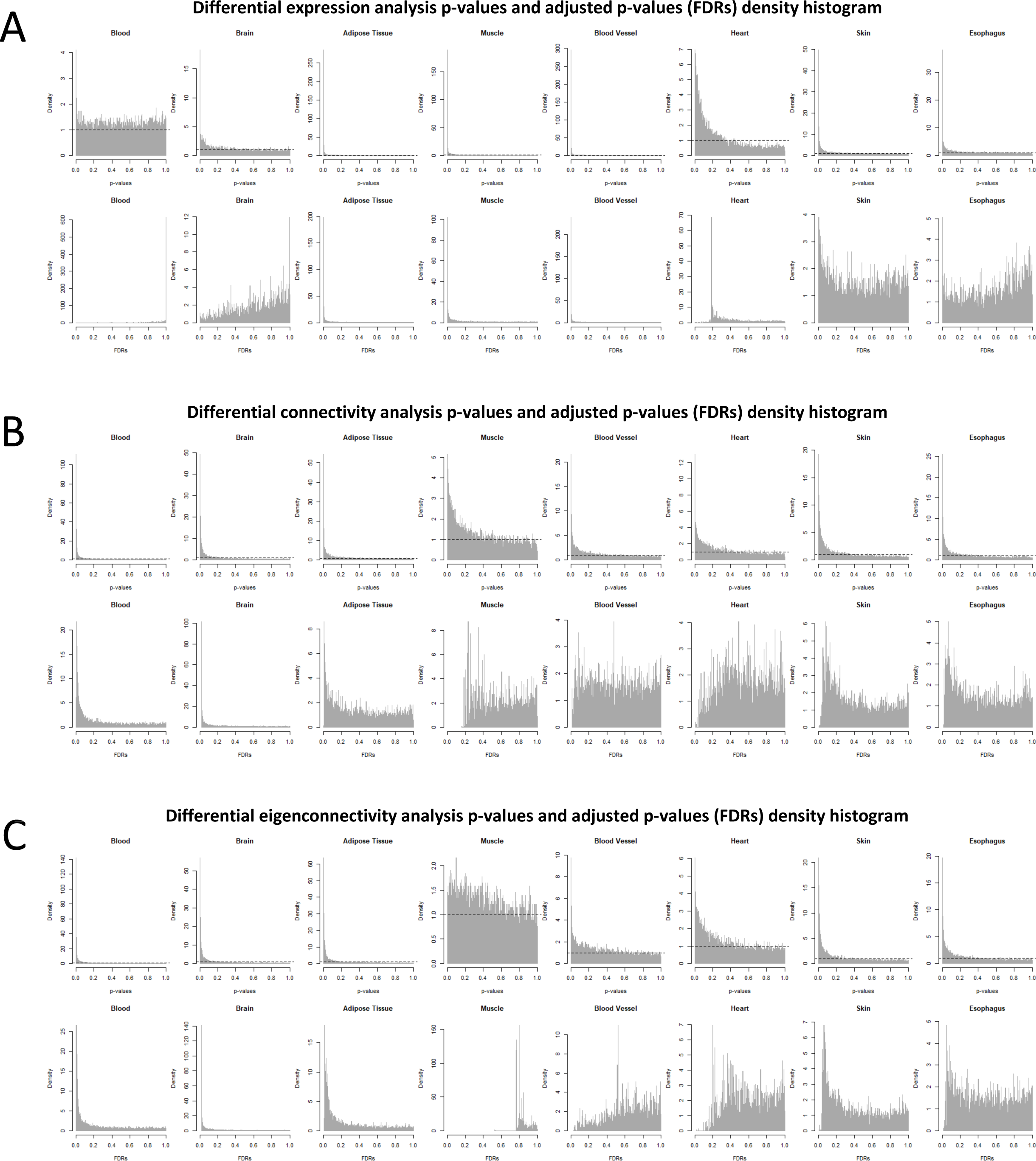
Density histogram of p-values and adjusted p-values (FDRs) of the regression analyses in each tissue. A) Distribution of p-values and adjusted p-values (FDRs) in the differential expression analysis. B) Distribution of p-values and adjusted p-values (FDRs) in the differential connectivity analysis. C) Distribution of p-values and adjusted p-values (FDRs) in the differential eigenconnectivity analysis. Dashed horizontal lines in the raw p-value density histograms represent what would be the expected distribution of p-values if the null hypothesis of no change was true for all genes.

**Figure S3.**
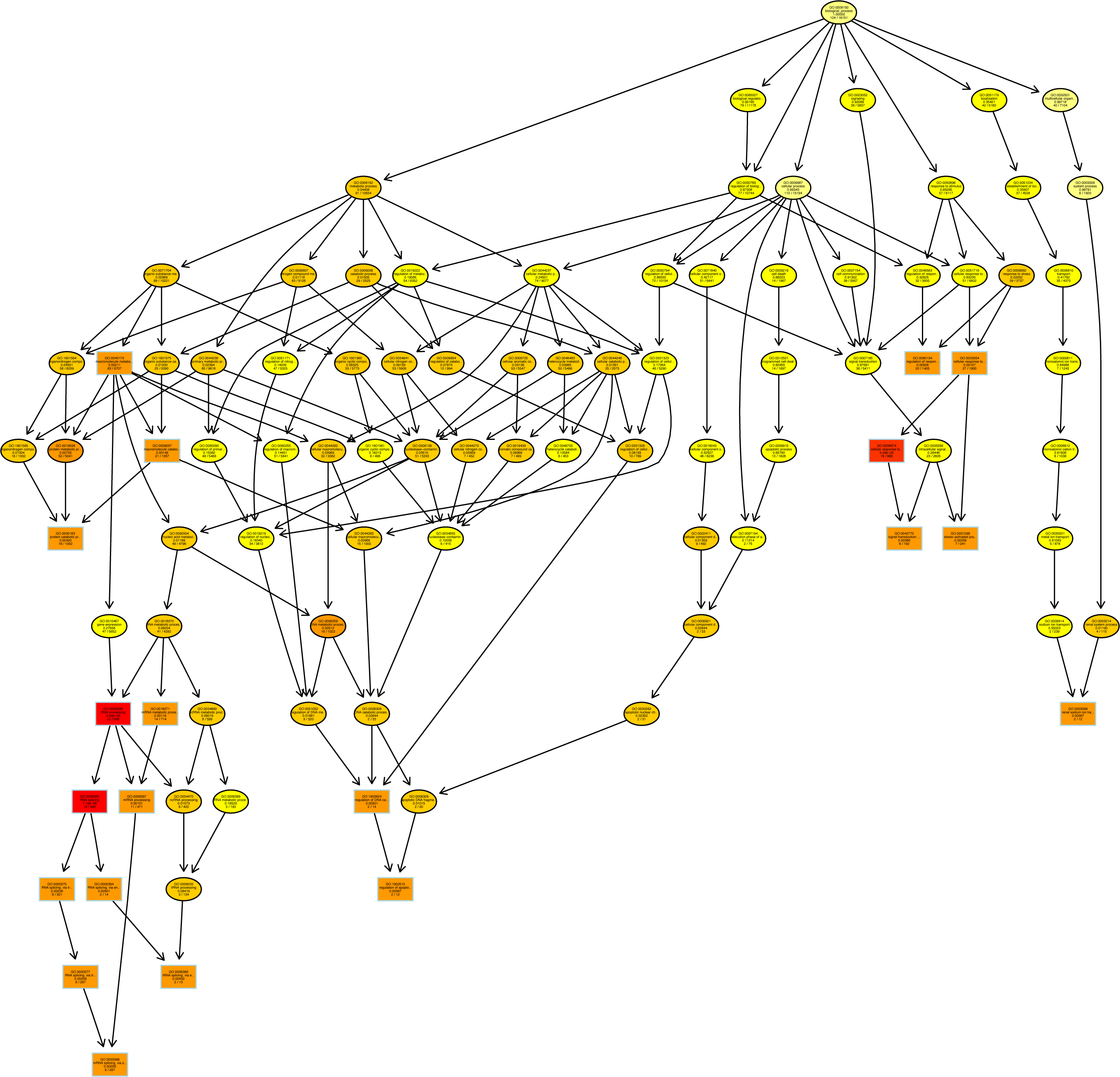
Induced GO subgraph of the 20 top enriched nodes in the 134-set.

**Figure S4.**
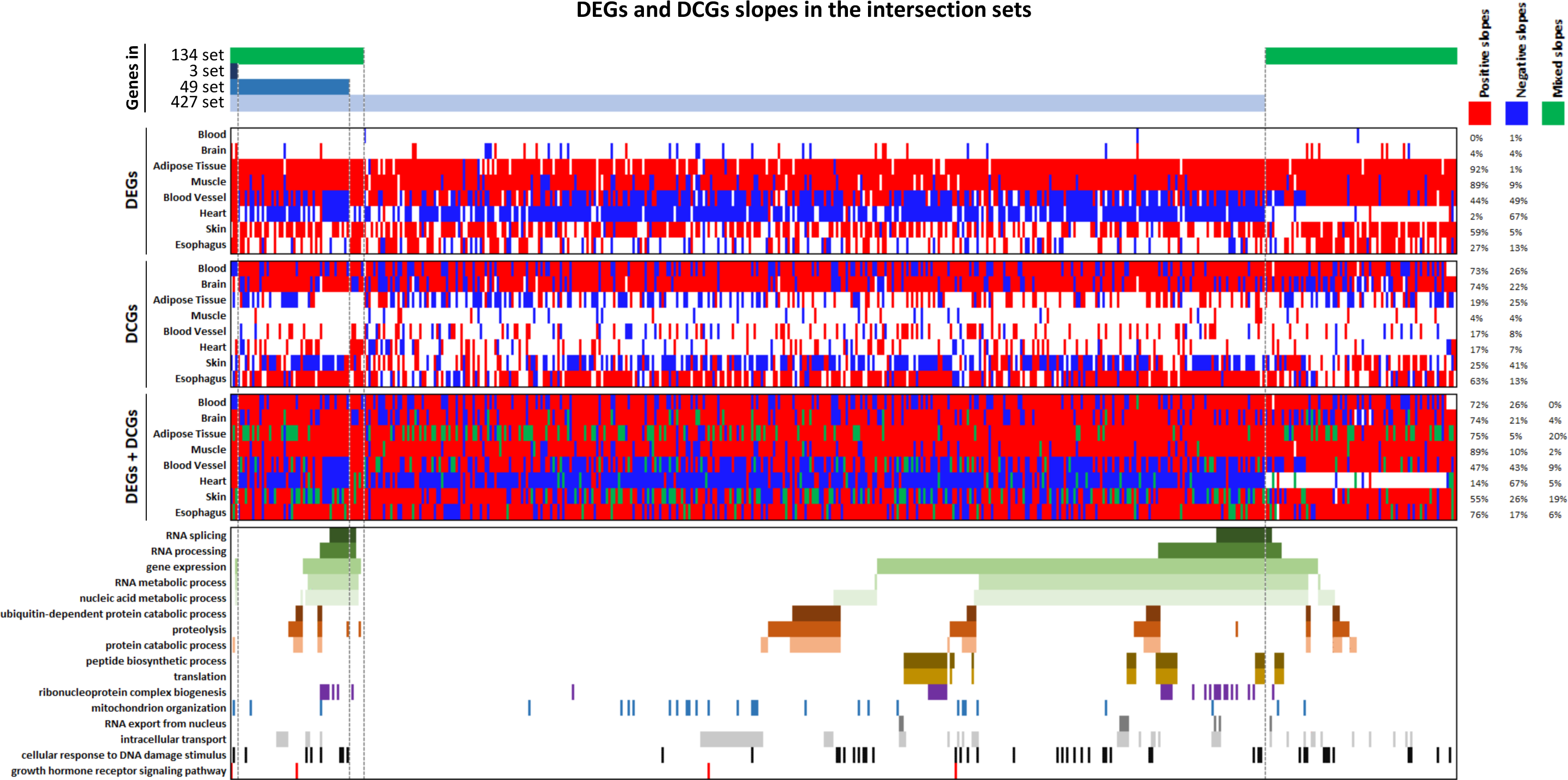
Slopes of DEGs and DCGs in the selected intersection sets. The most commonly annotated terms were selected and displayed in the bottom panel. In the two upper heatmaps, red entries represent positive slopes and blue entries represent negative slopes. In the bottom heatmap, red entries represent genes that are either a positive DEG, a positive DCG, or positive in both regressions; blue entries represent genes that are either a negative DEG, a negative DCG, or negative in both regressions; and green entries indicate genes significant in both regressions, but the slopes have opposite signals (either a positive DEG and negative DCG or a negative DEG and positive DCG).

**Figure S5.**
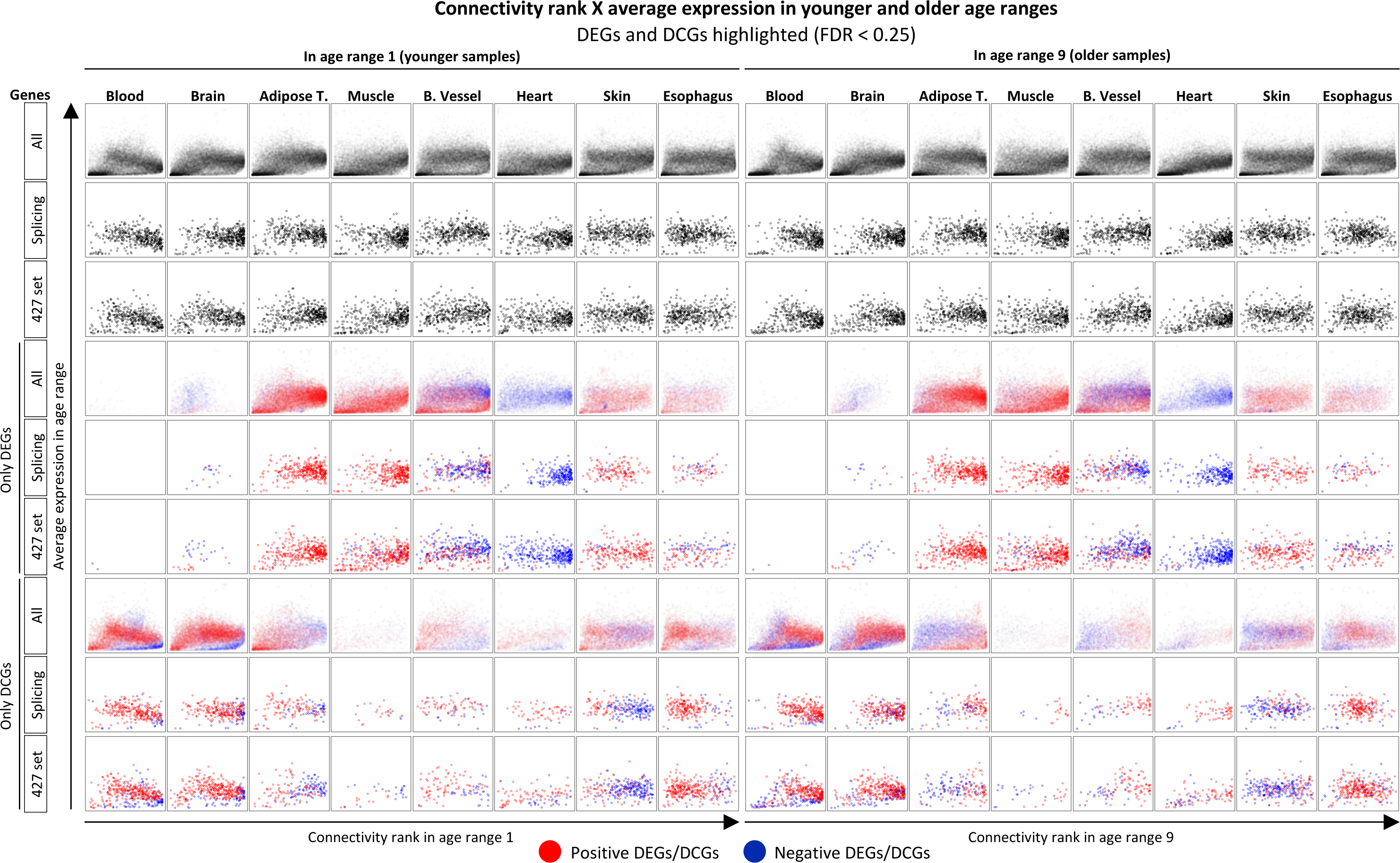
Connectivity ranks X average expression of different sets of genes in age ranges 1 and 9, with DEGs and DCGs highlighted. DEGs and DCGs highlighted considered FDR < 0.25, to emphasize the overall patterns of the trajectories. Results with FDR < 0.1 can be found in Supplemental Figure S6.

**Figure S6.**
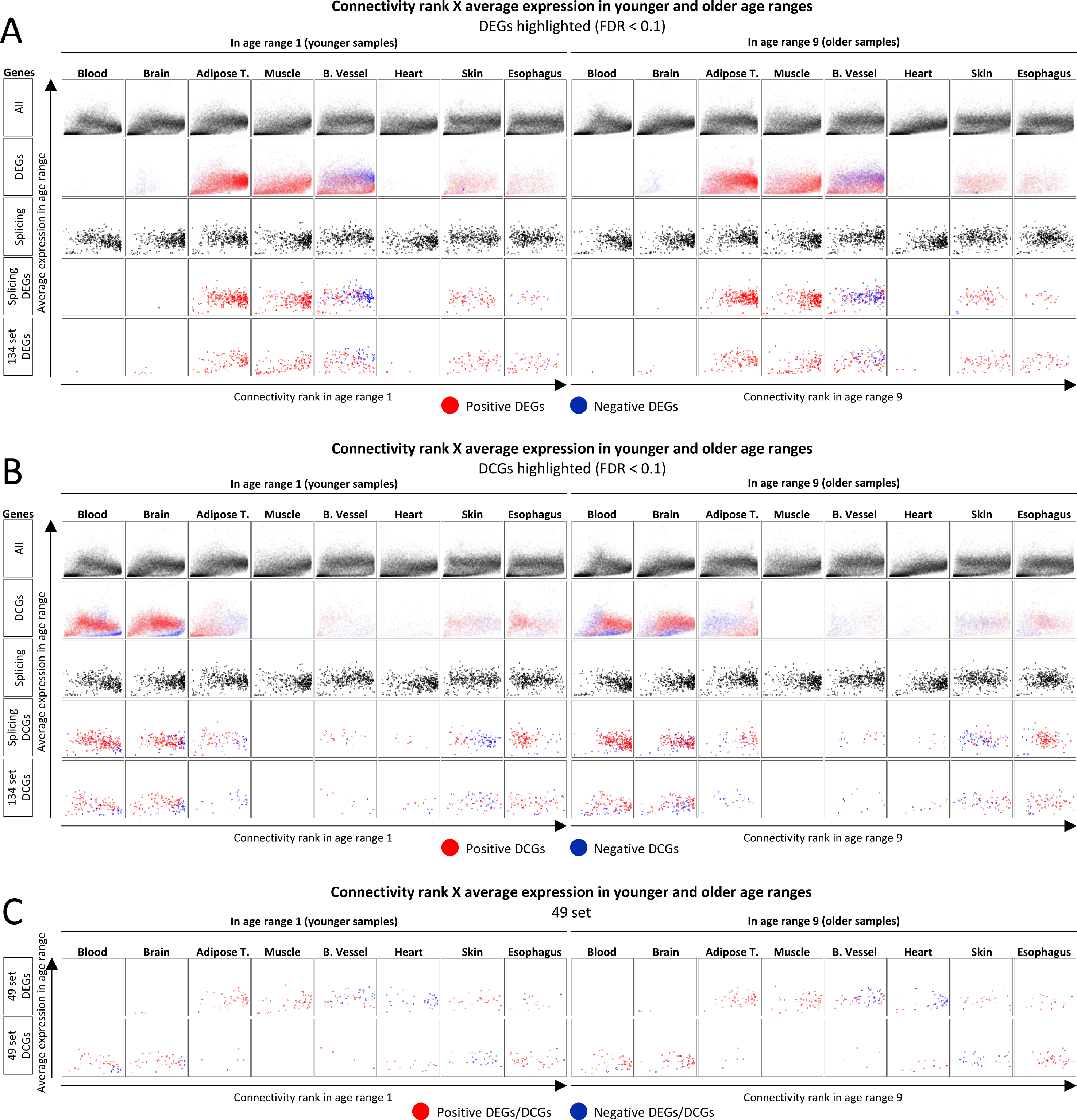
Distribution of positive and negative slopes among DEGs and DCGs. A) Connectivity ranks X average expression of different sets of genes in age range 1 and age range 9, with DEGs highlighted with FDR < 0.1. B) Connectivity ranks X average expression of different sets of genes in age range 1 and age range 9, with DCGs highlighted with FDR < 0.1. C) Connectivity ranks X average expression of the 49-set genes in age range 1 and age range 9, with DEGs and DCGs highlighted with FDR < 0.1 for all tissues except heart, which used an FDR < 0.25. Figure S6C only contains the charts for the 49-set because the other charts for the cases when other tissues are FDR < 0.1 or FDR < 0.25 are already included in Figures S5, S6A and S6B.

**Figure S7.**
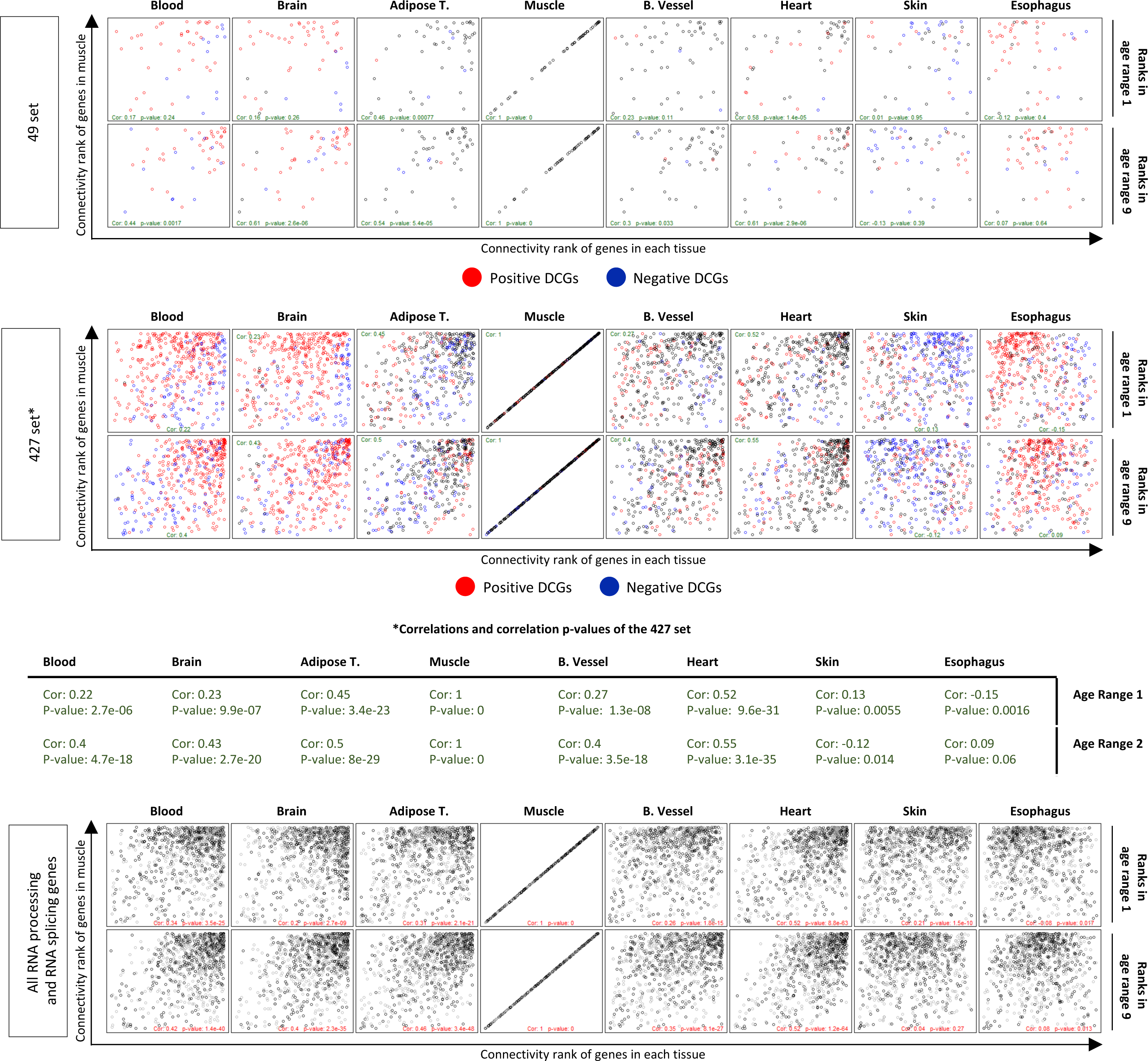
Connectivity ranks of genes in each tissue X connectivity rank of genes in muscle. The connectivity rank of genes in each tissue were plotted against the connectivity rank of genes in muscle in order to observe if there is convergence of the connectivity ranks with aging. Muscle was chosen as a reference because it does not have any DCG at FDR < 0.1. Plots presented refer to the 49-set, 427-set and all RNA processing and RNA splicing genes. Correlation and correlation p-values for the 427-set plot are displayed in the table under it due to lack of space in the chart.

**Figure S8.**
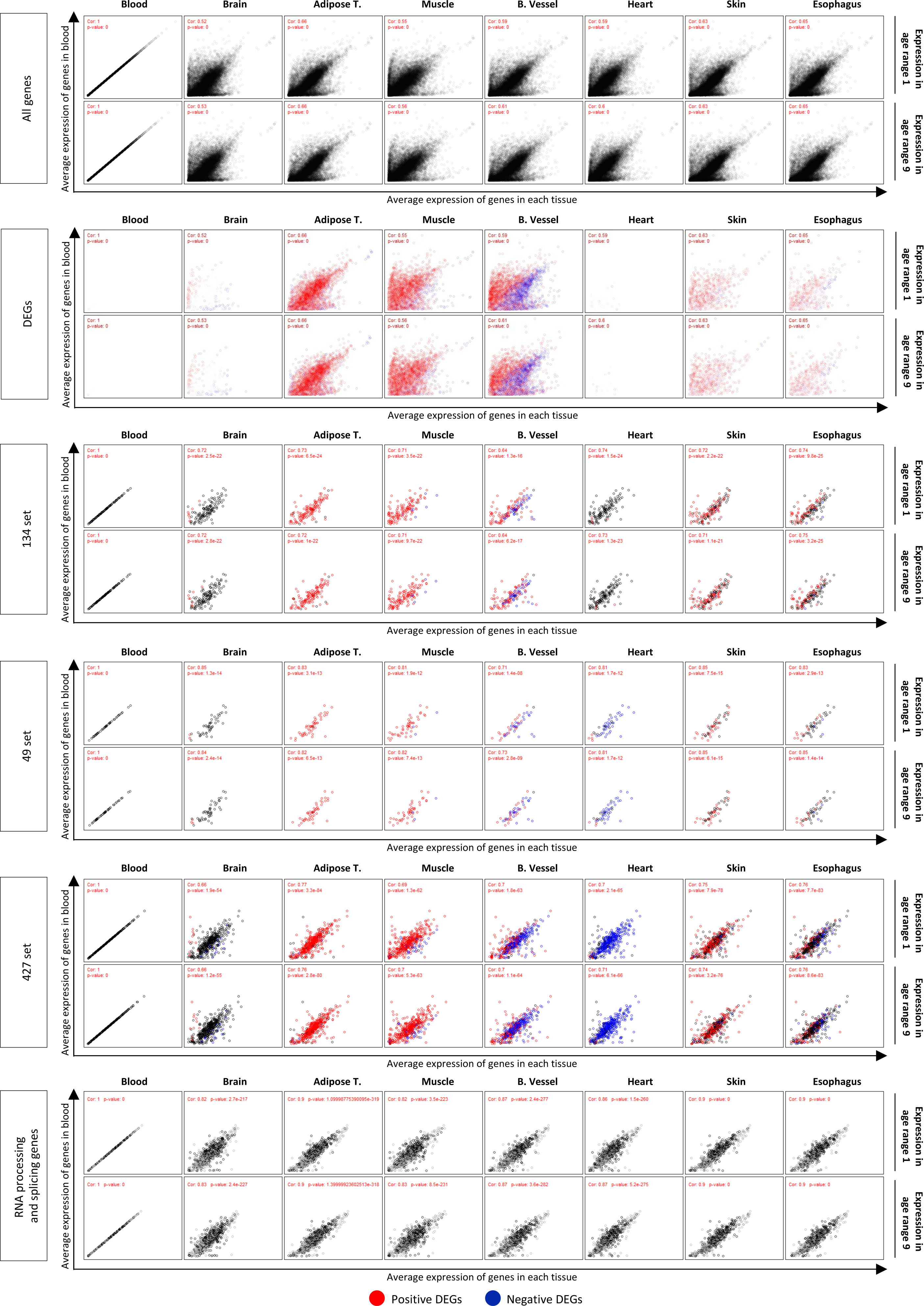
Average expression of genes in each tissue X average expression of genes in blood. The average expression of genes in each tissue was plotted against the average expression of genes in blood in order to observe if there is convergence of the expression with aging. Blood was chosen as a reference because it has a minimal amount of DEGs at FDR < 0.1. Plots presented refer to all genes, all DEGs, the 134-set, the 49-set, 427-set and all RNA processing and RNA splicing genes.

**Figure S9.**
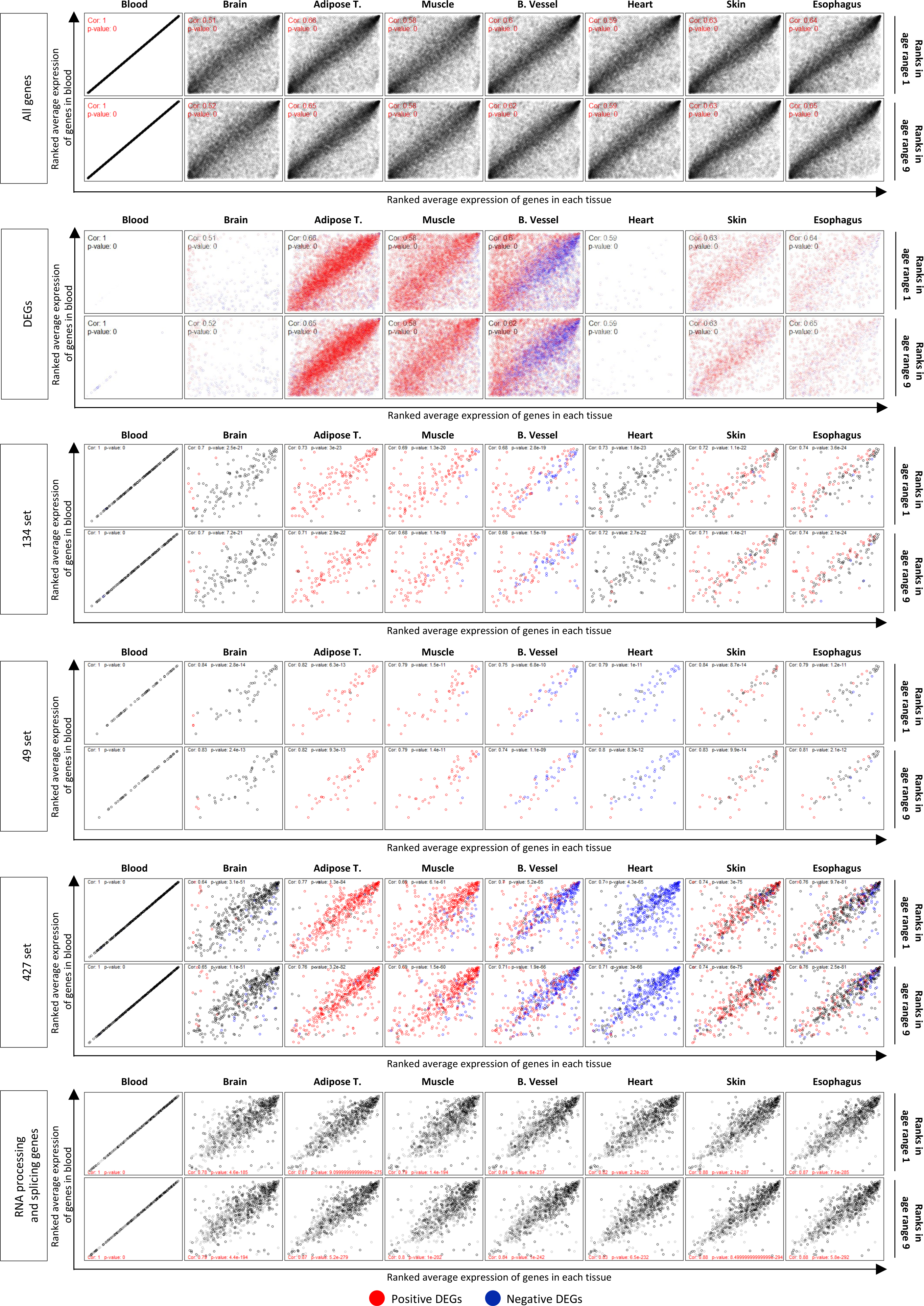
Ranked average expression of genes in each tissue X ranked average expression of genes in blood. The ranked average expression of genes in each tissue was plotted against the ranked average expression of genes in blood in order to observe if there is convergence of the expression with aging. Blood was chosen as a reference because it has a minimal amount of DEGs at FDR < 0.1. Plots presented refer to all genes, all DEGs, the 134-set, the 49-set, 427-set and all RNA processing and RNA splicing genes.

**Figure S10.**
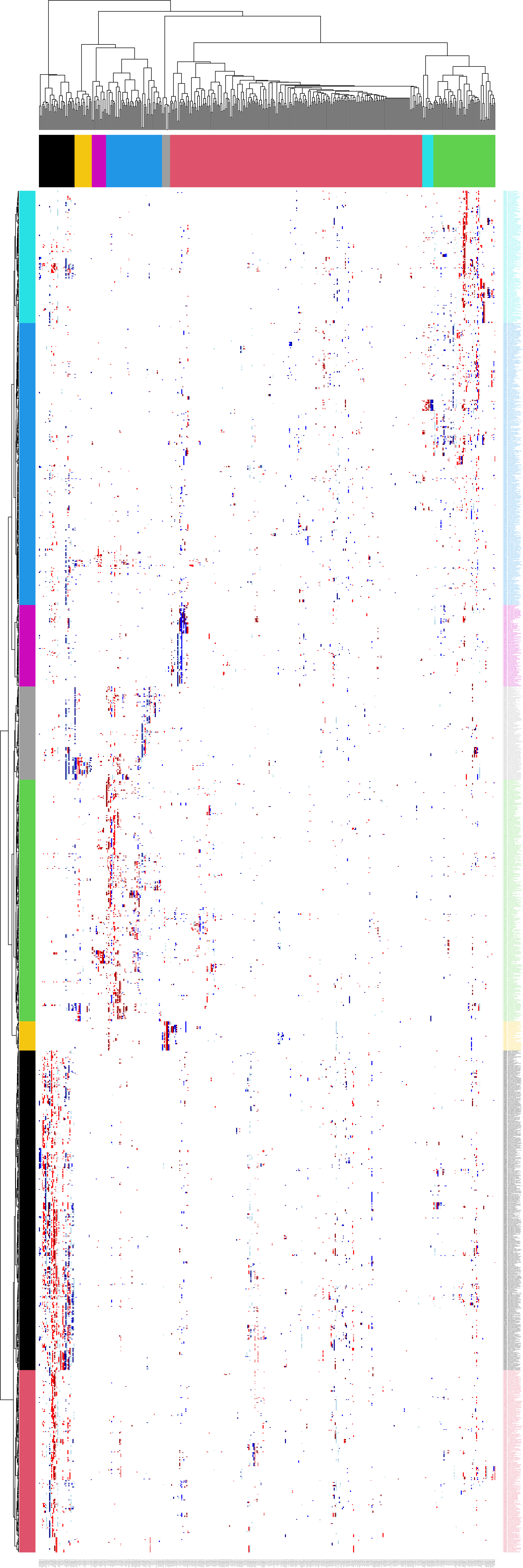
Consensus Modules Enrichment Heatmap. Larger version of the heatmap in Figure 5A, displaying all labels.

